# Regeneration of actin filament branches from the same Arp2/3 complex

**DOI:** 10.1101/2023.09.22.558980

**Authors:** Foad Ghasemi, LuYan Cao, Miroslav Mladenov, Bérengère Guichard, Michael Way, Antoine Jégou, Guillaume Romet-Lemonne

## Abstract

Branched actin filaments are found in many key cellular structures. Branches are nucleated by the Arp2/3 complex activated by nucleation-promoting factor (NPF) proteins and bound to the side of pre-existing ‘mother’ filaments. Over time, branches dissociate from their mother filament, leading to network reorganization and turnover, but this mechanism is less understood. Here, using microfluidics and purified proteins, we examined the dissociation of individual branches under controlled biochemical and mechanical conditions. We observe that Arp2/3 remains bound to the mother filament after most debranching events, even when accelerated by force. Unexpectedly, this mother-remaining Arp2/3 readily nucleates a new actin filament branch, without being activated anew by an NPF: it simply needs to exchange its nucleotide and bind an actin monomer. The protein GMF, which accelerates debranching, prevents branch re-nucleation. Our results suggest that actin filament re-nucleation can provide a self-repair mechanism, helping branched networks to sustain mechanical stress in cells over extended periods of time.

## INTRODUCTION

In cells, branched actin filaments are involved in a variety of essential processes. They are key components of the cell cortex, of the lamellipodium at the front of a migrating cell, and of endocytic cups (*1, 2*). Branched actin filaments are also important to displace and reshape mitochondria in the cytoplasm (*3*), and to repair damaged chromatin in the nucleus (*4*). These filament networks have specific dynamics, as both their assembly and disassembly are precisely controlled. In most of these processes, branched actin networks play a mechanical role, exerting forces as the filaments elongate, and adapting to mechanical loads (*5*–*7*).

An actin filament branch is nucleated by the Arp2/3 complex bound to the side of a pre-existing “mother” filament (reviewed in (*8*)). The Arp2/3 complex is composed of seven subunits, including Actin-related proteins (Arp) 2 and 3 which can template the polymerization of an actin filament. In its basal state, however, the Arp2/3 complex adopts an inactive conformation where Arp2 and Arp3 are splayed apart and adopt a twisted conformation similar to monomeric actin (G-actin), and thus cannot nucleate a filament (F-actin) (*9*–*13*). Activation is mediated by membrane-anchored nucleation promoting factors (NPFs), whose VCA domains simultaneously bind Arp2/3 and G-actin. This results in a conformational change of the Arp2/3 complex, allowing it to bind to the side of the mother filament and to reposition Arp2 and Arp3 into a filament-like conformation. Upon detachment of the VCA domains, the new barbed end will elongate and form a branch, or “daughter” filament, with a well-characterized 70° angle with respect to the mother filament (*14*).

Like actin, Arp2 and Arp3 have a nucleotide-binding pocket. The nucleation of the branch requires the Arp2/3 complex to be loaded with ATP, which will be hydrolyzed after the branch has nucleated (*15*–*18*). As long as it remains in the branch junction, the Arp2/3 complex is not able to exchange its nucleotide, which will thus be a marker of its age: older branches will have an ADP-Arp2/3 at their junction.

Branches eventually dissociate from the mother filament, thereby contributing to the reorganization and the turnover of the branched network. In spite of its importance for the control of actin turnover, the mechanism of branch dissociation is less understood than that of branch nucleation (*19, 20*). In vitro studies have shown that ATP hydrolysis in the Arp2/3 complex favors branch dissociation: “older” ADP-Arp2/3 branches dissociate more easily (*17, 19, 21*).

Further, the Arp2/3 complex can be directly targeted by the protein GMF to accelerate debranching, or by the protein cortactin to stabilize the branch (*21*–*25*). In addition, Pandit et al. have recently shown that the application of sub-piconewton pulling forces to branches dramatically accelerates their dissociation (*21*). This observation is of prime importance because branched actin networks are exposed to mechanical forces in cells, and this raises the question of how they are able to sustain prolonged mechanical stress. It also means that one needs to control the mechanical context in order to properly study branch dissociation.

In spite of these recent advances, basic aspects of branch dissociation are still unclear. For instance, whether the Arp2/3 complex remains on the mother filament or departs with the dissociated branch remains elusive (*26*). Nonetheless, it is often considered that branch dissociation results from the detachment of the Arp2/3 complex from the mother filament (*14, 21*). In this scenario, the pointed end of the dissociated branch could remain capped by Arp2/3, preventing depolymerization and reannealling with a free barbed end. Recent cryo-EM data of branch junctions provide accurate descriptions of buried surface areas between the Arp2/3 complex and the mother and daughter filaments (*27*–*29*). Yet, they do not provide estimations of the free energy associated with these interfaces, and their relative strengths remains an open question.

To address this question, we have monitored the dissociation of branches in controlled mechanical conditions. We find that, not only does Arp2/3 remain bound to the mother filament after nearly all debranching events, but, strikingly, it can then rapidly reload fresh ATP and nucleate a new branch, without being reactivated by NPFs.

## RESULTS

All experiments were carried out *in vitro* at 25°C, using purified proteins (from mammalian origin unless specified otherwise), in a buffer at pH 7.0 containing 50 mM KCl (see Methods). In most experiments, we used alpha-skeletal actin, but we also used gamma-cytoplasmic actin where specified.

### Branch dissociation and re-nucleation

We used a microfluidics assay (*30*–*32*) to monitor the nucleation and the dissociation of branches while exposing them to controlled mechanical tension (Fig. 1A). We first elongated filaments from surface-anchored seeds by flowing fluorescently labeled actin monomers (G-actin) in the microchamber. We subsequently generated branches by flowing in Arp2/3 together with the VCA region of the NPF N-WASP and G-actin labeled with a different fluorescent color in order to facilitate the identification of branches, which tend to align with the mother filament because of the flow (see Methods). We next flowed labeled G-actin alone in the microchamber, in order to monitor the elongation and the dissociation of the branches without nucleating new branches. This allowed us to keep the branch density low enough to clearly monitor the fate of individual branches (Fig. 1B,C).

**Fig. 1.**
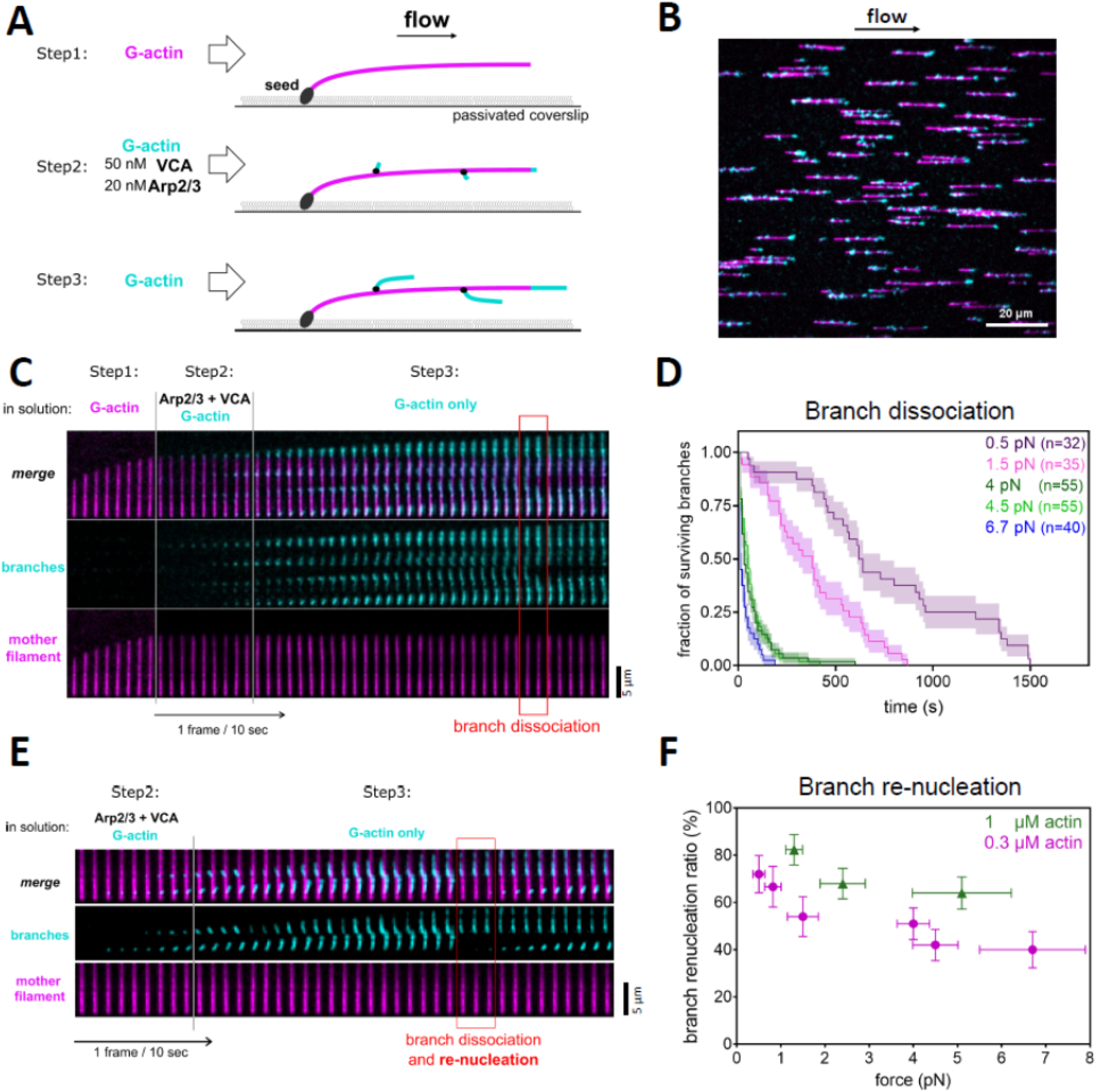
New actin branches can grow from the same Arp2/3 complex after a branch dissociation event. (**A**) Schematic of a typical branching and debranching experiment using microfluidics. Filaments are first elongated from surface-anchored spectrin-actin seeds by flowing in Alexa488(10%)-G-actin (magenta). Branches are then nucleated by flowing in Alexa568(10%)-G-actin (cyan) with VCA and Arp2/3. They are further elongated by flowing in Alexa568(10%)-G-actin alone. The elongation and dissociation of branches can then be monitored over time. The flowing solution applies a pulling force to the mother filament and the branches. (**B**) Portion of a TIRF microscopy field of view, showing mother filaments (magenta) and branches (cyan) formed as indicated in (A). (**C**) Time lapse showing the elongation of a mother filament (magenta), the nucleation of branches (cyan), their elongation, and the dissociation of a branch (red box). (**D**) Pulling forces accelerate the dissociation of branches. Each curve shows the surviving fraction of n branches monitored over time, for a different experiment (branches aged for 20 minutes with 0.3 µM actin, between step 2 and 3 indicated in panel (A)). Each experiment was carried out with a different flow rate, in order to apply different forces, indicated as the average force applied to the branches as they dissociated. (**E**) Timelapse, similar to the one in (C), showing an example of branch re-nucleation (red box). (**F**) Pulling forces moderately reduce the fraction of branches that re-nucleate after dissociation. Each point is from a single experiment, analyzing, from left to right, n=32, 30, 35, 55, 55, 40 branches (0.3 µM actin) and n=34, 50, 50 branches (1 µM actin). Error bars are standard deviations.

We quantified the dissociation of branches over time, as we exposed them to different flow velocities and thus to different forces (see Methods and (*31, 32*)). Similar to Pandit et al. (*21*), we observed that branches dissociated faster from mother filaments when exposed to increasing forces in the picoNewton range (Fig. 1D).

Unexpectedly, however, we observed that the majority of dissociation events were followed by the growth of a new branch from the same location (Fig. 1E, Supp Fig S1). The regeneration of the branches took place immediately after branch dissociation, with no detectable lag (Supp Fig S2). The fraction of branches that regenerated after dissociating, a percentage we call the “branch re-nucleation ratio”, decreased with tension, but remained above 50% for forces up to 5 pN (Fig 1F). We also observed that the branch re-nucleation fraction increased with G-actin concentration (Fig 1F). The branch re-nucleation events took place in the absence of Arp2/3 and VCA in solution, implying that Arp2/3 remained on the mother filament upon dissociation of the branch, and that a new branch could be nucleated from that same Arp2/3.

Actin filaments rarely fragment in our assay, in the absence of specific severing proteins such as ADF/cofilin (*32, 33*). Nevertheless, any severing event occurring close to the branching junction would be mistaken for a dissociation event, and the subsequent elongation of the severed branch would be wrongly interpreted as re-nucleation. We thus sought to determine whether our observation of branch dissociation events was contaminated by branch severing events. First, we verified that the branch re-nucleation ratio was unaffected by changes in actin fluorescent labeling and illumination (Supp Fig S3A). Second, we designed a specific experiment to test if any actin subunits from the first nucleation event were still present in the re-nucleated branch (Supp Fig S3B-D). This experiment is similar to Fig 1A, except that we used G-actin with a high labeling fraction of a third fluorophore for the branching reaction (step 2), resulting in branches with a clearly labeled pointed end. After branch dissociation and the nucleation of new branches, these fluorescently-labeled actin subunits could no longer be detected at the branch junction, in illumination conditions where single fluorophores could be detected. We conclude that the branch departure events that we observe are genuine branch dissociation events, where no actin subunits from the initial daughter filament remain bound to the Arp2/3 complex associated to the mother filament.

Our observation that Arp2/3 remained bound to the mother filament upon dissociation of the branch appeared to contradict Pandit et al. who detected no fluorescently labeled Arp2/3 on the mother immediately after branch dissociation (*21*). We wondered if this difference could come from our using mammalian Arp2/3 and mammalian actin, while Pandit et al. used Arp2/3 from fission yeast and mammalian actin. To test this hypothesis, we repeated our experiments with Arp2/3 from budding yeast, and observed a branch re-nucleation ratio of less than 10% (Supp Fig S4B). Our interpretation is that the interface between the mother filament and the Arp2/3 complex is much weaker when the latter is from yeast while the former is made of mammalian actin. Consistently, we find that branches dissociate much faster when using yeast Arp2/3 instead of mammalian Arp2/3 (Supp Fig S4A).

### Impact of force orientation

In our standard experiment, the mother filaments aligned with the flow, and thus the pulling force applied to the branches is parallel to the mother filaments (Fig 1). We wondered if the orientation of the force impacts the branch dissociation rate and re-nucleation. In order to vary the angle between the mother filament and the flow, we modified our assay as follows (Fig 2, Methods).

**Fig. 2.**
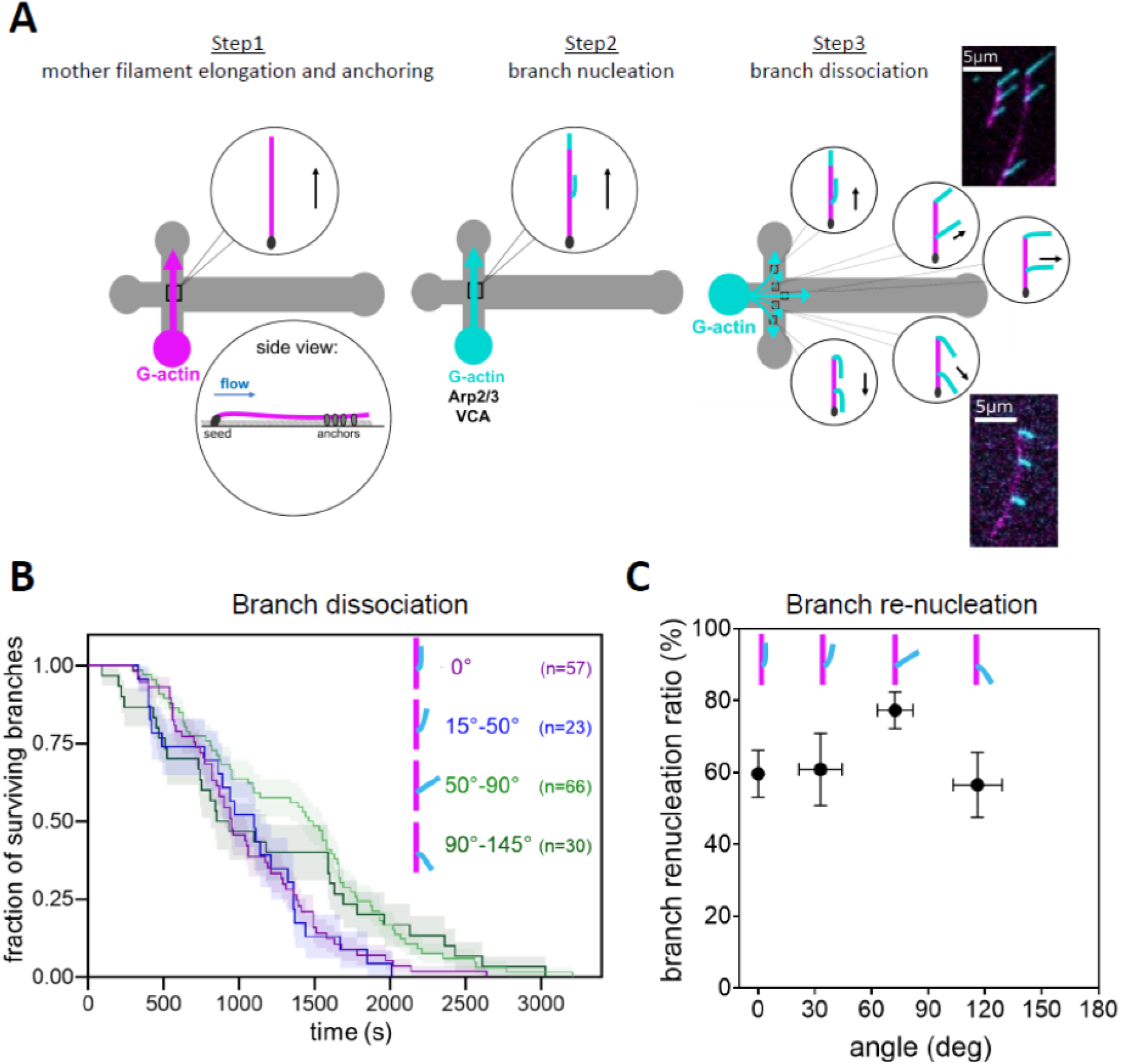
Impact of force orientation on debranching and re-nucleation. (**A**) Schematic of the assay where branches are pulled on with different orientations. Mother filaments (magenta) are elongated with a biotinylated segment which is anchored to the surface (step 1). During the debranching and re-nucleation phase (step 3), the pulling force follows the direction of the flow, which makes different angles with respect to the mother filaments in different regions of the microfluidics chamber. The microscope images show examples from two different fields, with forces applied on branches at 30-47° (top image) and at 114-120° (bottom image) with respect to their mother filaments. (**B**) Each curve shows the surviving fraction of branches monitored over time, for a different population of n branches from 9 independent experiments pooled together according to the angle between the applied force and the mother filament. (**C**) The direction of the pulling force has a moderate impact on the branch re-nucleation ratio, computed for each of the four populations of branches whose detachment is shown in (B). The value for 50-90° is significantly higher than for smaller angles (p=0.033, two-sided Fisher’s exact test). Error bars indicate binomial standard deviations.

We worked in the region of the microchamber where the channels intersect, so we can orient the flow of the incoming solutions in different directions. We grew filaments from surface-anchored seeds by flowing in fluorescently labeled G-actin, as before, but we then elongated them further with a solution supplemented with biotinylated G-actin. This created a biotinylated filament segment, that we could then anchor to surface-anchored biotin-BSA, by flowing in neutravidin. As a result, the orientation of mother filaments was fixed, imposed by the anchored seed and the anchored biotinylated segment, regardless of the orientation of the flow. We then nucleated and elongated branches, and monitored their fate as before. To avoid potential artifacts due to local anchoring points, we only followed branches whose junction was in the unanchored segment of the mother filament.

By flowing the G-actin solution from one channel into the three others, we obtained different local flow directions, so that applied forces have different angles with respect to the mother filaments, within the same microchamber (Fig 2A). Mother filaments whose biotinylated segment failed to bind the surface provided local control situations where the mother and the branch were aligned with the flow. In a separate experiment, we verified that the local flow velocity was the same for the different flow directions, by tracking the movement of micrometer-size beads (Supp Fig S5). The forces we applied to the actin filament branches thus had different orientations but similar amplitudes (approximately 1 pN).

We pooled branches into four subpopulations, based on the angle θ between the local direction of the flow (i.e. the direction of the applied force) and the mother filament. We found that branches dissociated at a similar rate from their mother filaments, regardless of the orientation of the force (Fig 2B). We found that the branch re-nucleation ratio was very mildly affected across the range of angles we studied (0-145°).The branch re-nucleation ratio was approximately 30% higher when θ was in the vicinity of 70°, the canonical angle of the branch junction (Fig 2C).

These results show that branch re-nucleation following debranching is a general mechanism, taking place regardless of the orientation of the pulling force. The higher branch re-nucleation ratio measured around θ = 70° indicates that Arp2/3 is even more likely to remain bound to the mother filament when the tension is applied without bending the branch and without shearing the branch junction.

### Actin isoform and profilin

Our results so far imply that, following the majority of branch dissociation events, Arp2/3 remains on the mother filament and nucleates a new branch. Since we made these observations using alpha-skeletal actin, we wondered whether they would differ if we used cytoplasmic actin. To be closer to physiological conditions, where actin is unlabeled, we modified our assay to monitor branch junctions made with unlabeled cytoplasmic gamma-actin (Fig 3A-B, Methods). Briefly, we successively flowed different actin solutions in the microchamber, to elongate mother filaments comprising a long segment made entirely of unlabeled gamma-actin, between two segments made of labeled alpha-actin from skeletal muscle. Next, we nucleated branches using unlabeled gamma-actin. Finally, we elongated the branches and monitored their outcome while continuously flowing labeled alpha-actin in the chamber. This enabled us to monitor the dissociation of branches where only unlabeled gamma-actin was in contact with the Arp2/3 complex, in both the mother filament and the pointed region of the branch. We found that the branch re-nucleation ratio was the same as for our standard branches, monitored simultaneously in the same microchamber (Fig 3C, Methods).

**Fig. 3.**
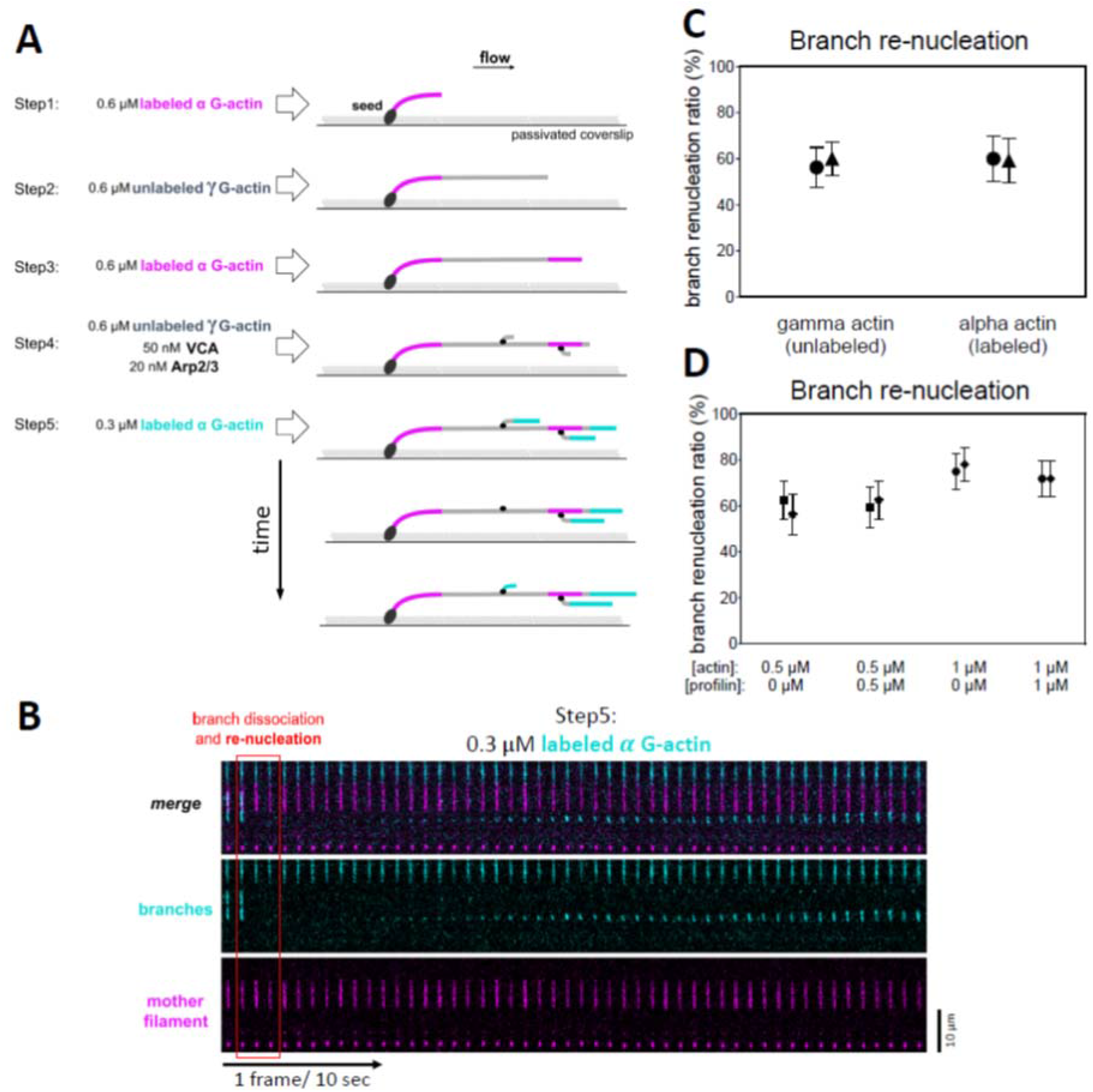
Branch re-nucleation is not actin isoform-specific and occurs in the presence of profilin. (**A**) Schematic of experiment using microfluidics to study branch junctions made with unlabeled cytoplasmic gamma-actin. The mother filaments comprise a long segment made with unlabeled cytoplasmic gamma-actin (steps 1-3), and the branches are nucleated using unlabeled cytoplasmic gamma-actin (step 4). (**B**) Timelapse showing a branch dissociating and re-nucleating, from the unlabeled segment of the mother filament, during step 5 of the experiment described in (A). (**C**) The branch re-nucleation ratio is similar for mother filaments made with unlabeled cytoplasmic gamma-actin or with Alexa488-labeled(10%) alpha-skeletal actin. (**D**) The branch re-nucleation ratio is not affected by adding equimolar amounts of profilin to G-actin during the final step of the experiment (branch dissociation and re-nucleation, step 3 in Fig 1A). In (C,D) each data point is from one experiment, monitoring n=32, 45, 25, 27 dissociating branches (from left to right) for experiments in (C) and n=32 dissociating branches for each experiment in (D). Points with matching symbols are from observations carried out simultaneously, in different regions of the same microfluidics chamber. Error bars indicate standard deviations.

We also wondered if our results could be specific to the NPF we used to nucleate the first branches, VCA from N-WASP. We thus repeated our experiments using VCA from WASP and found that 55% (±11%, n=22) of branches were regenerated after dissociating from the mother filament. This is similar to what we observed for branches nucleated using VCA from N-WASP.

Since, in cells, available actin monomers are mostly in complex with profilin (*34*), we asked whether profilin could affect branch re-nucleation. We compared, side-by-side in the same microchamber, the behavior of branches in the presence of actin alone and supplemented with profilin (Fig. 3D). We found that adding equimolar amounts of profilin to actin had no impact on the branch re-nucleation ratio.

### Nucleotide exchange on mother-remaining Arp2/3

Having established that branch regeneration is a robust mechanism, we next sought to better understand the underlying molecular mechanism.

Earlier reports indicate that Arp2/3 needs to be loaded with ATP in order to be activated by an NPF and nucleate a branch (*15, 16*). This led us to wonder whether this was also the case for the re-nucleation events we observed. In order to determine if branches were re-nucleated from ADP-or ATP-Arp2/3, we took advantage of the fact that ADP-Arp2/3 branches dissociate faster (*17, 19, 21*).

To obtain ADP-Arp2/3 branches, we aged them for 20 minutes before monitoring their dissociation (Fig 4A). As expected, in our experiments, the surviving fraction of aged branches (i.e. mostly ADP-Arp2/3 at the start of the experiment) decreased faster than that of non-aged branches (i.e. ATP-Arp2/3 and ADP-Pi-Arp2/3 at the start of the experiment) (Fig 4A, B).

**Fig. 4.**
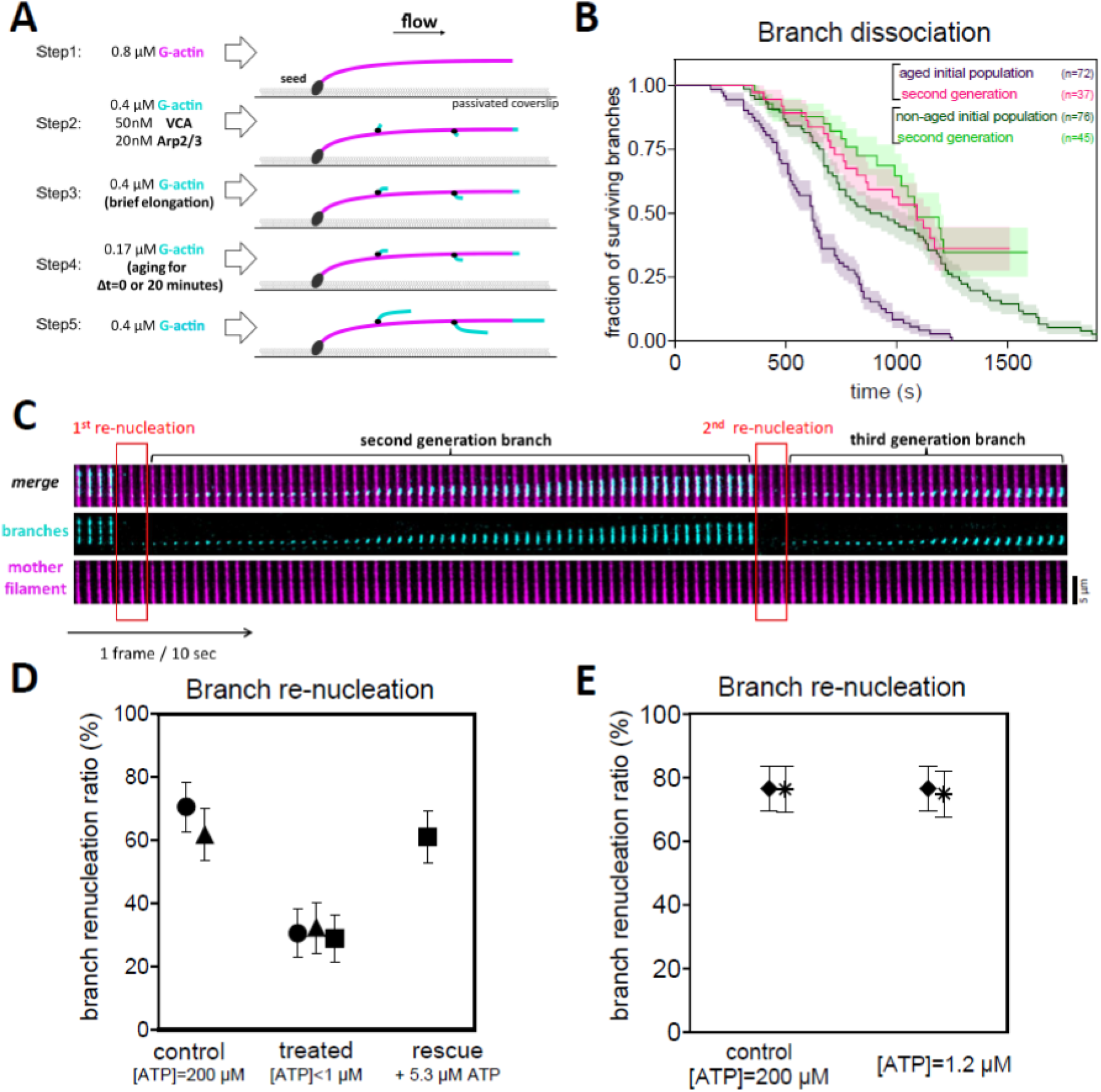
Arp2/3 binds fresh ATP to re-nucleate a branch. (**A**) Schematic of the aging experiment using microfluidics. Compared to the standard assay shown in Fig 1, an extra step (step 4) is added in order to age the branch junctions without elongating the branches: a lower concentration of G-actin is flowed in for 0 min (non-aged branches) or 20 min (aged branches). (**B**) Kinetics of branch dissociation, for aged (purple curve) and non-aged branches (dark green curve), and for the branches that have re-nucleated in each case (“second generation”). (**C**) Time-lapse showing a branch detaching and re-nucleating, twice. In this experiment, 71% of the initial branches (n=41) re-nucleated, and 69% of the second generation of branches (n=29) re-nucleated. (**D**) The branch re-nucleation ratio decreases when ATP is depleted from the buffer, and is rescued by adding back ATP. (**E**) The Branch re-nucleation ratio is not affected when ATP is diluted to 1.2 µM. In (D,E) each data point is from one experiment, monitoring (from left to right) n=34, 34, 36, 34, 38, 36 dissociating branches for experiments in (D) and n=34, 30, 35, 30 dissociating branches for each experiment in (E). Points with matching symbols are from observations carried out simultaneously, in different regions of the same microfluidics chamber. Error bars indicate standard deviations.

Strikingly, in both situations we observed that the re-nucleated branches dissociated slowly, like non-aged branches (Fig 4B). This suggests that Arp2/3 is loaded with ATP when it re-nucleates a filament, after dissociation of the first branch.

The fact that aged branches dissociate faster than non-aged branches (Fig 4B) implies that most mother-remaining Arp2/3 complexes are in the ADP-state. Thus, our result that branches are re-nucleated from ATP-Arp2/3 indicates that Arp2/3 exchanges its nucleotide after branch dissociation, in order to nucleate a new branch. Consistently with this, we observed that re-nucleated branches, after they dissociated, were re-nucleated again, with the same branch re-nucleation ratio (Fig 4C). This could be observed for multiple generations, as if each re-nucleation event was independent of the past history of that specific Arp2/3 complex.

To further confirm the notion that the mother-remaining Arp2/3 exchanges its nucleotide in order to nucleate a new branch, we decreased the ATP concentration to see if that would impair branch re-nucleation. To maintain G-actin in the ATP state, we did not fully remove ATP from solution. We verified, by measuring the elongation rate at filament barbed ends, that all the G-actin was indeed loaded with ATP in our experiments, even at our lowest concentrations of ATP. We observed that branches dissociating while exposed to G-actin in an ATP-deficient solution ([ATP]<1 µM, see methods) re-nucleated more than 2-fold less frequently than in our control experiment ([ATP]=200 µM) carried out simultaneously in the same microchamber (Fig 4D).

This result confirms that ATP in solution is required for branch re-nucleation. We were not able to accurately measure ATP concentrations below 1 µM, and thus could not determine its affinity for Arp2/3 bound to the mother filament. However, we measured that branch re-nucleation was unaffected when the ATP concentration was decreased as low as 1.2 µM (Fig 4E).

Together, our results indicate that actin filament branches dissociate from the ADP-Arp2/3 complex which remains on the mother filament, and that this mother-remaining Arp2/3 complex is able to rapidly exchange its nucleotide to become ATP-Arp2/3 and re-nucleate a branch.

### Detachment of mother-remaining Arp2/3

In order to estimate the rate at which the mother-remaining Arp2/3 exchanges its nucleotide, we needed to take into account competing reactions. We reasoned that, following branch dissociation, the mother-remaining ADP-Arp2/3 can either exchange its nucleotide to become ATP-Arp2/3, or dissociate from the mother filament. In turn, the ATP-Arp2/3 complex can either dissociate from the mother filament, or bind an actin monomer, which we assumed would stabilize its interaction with the mother filament as in a new branch junction. This last assumption is consistent with our observation that the branch re-nucleation ratio increases with the G-actin concentration (Fig 1F).

We thus sought to quantify the dissociation of the Arp2/3 complex from the mother filament. To do so, we fluorescently labeled the Arp2/3 complex (Methods) and verified that this labeling did not affect the nucleation, dissociation and re-nucleation of the branches (Fig 5A). We could thus directly visualize the same Arp2/3 complex remaining bound to the mother filament throughout branch dissociation and re-nucleation.

**Fig. 5.**
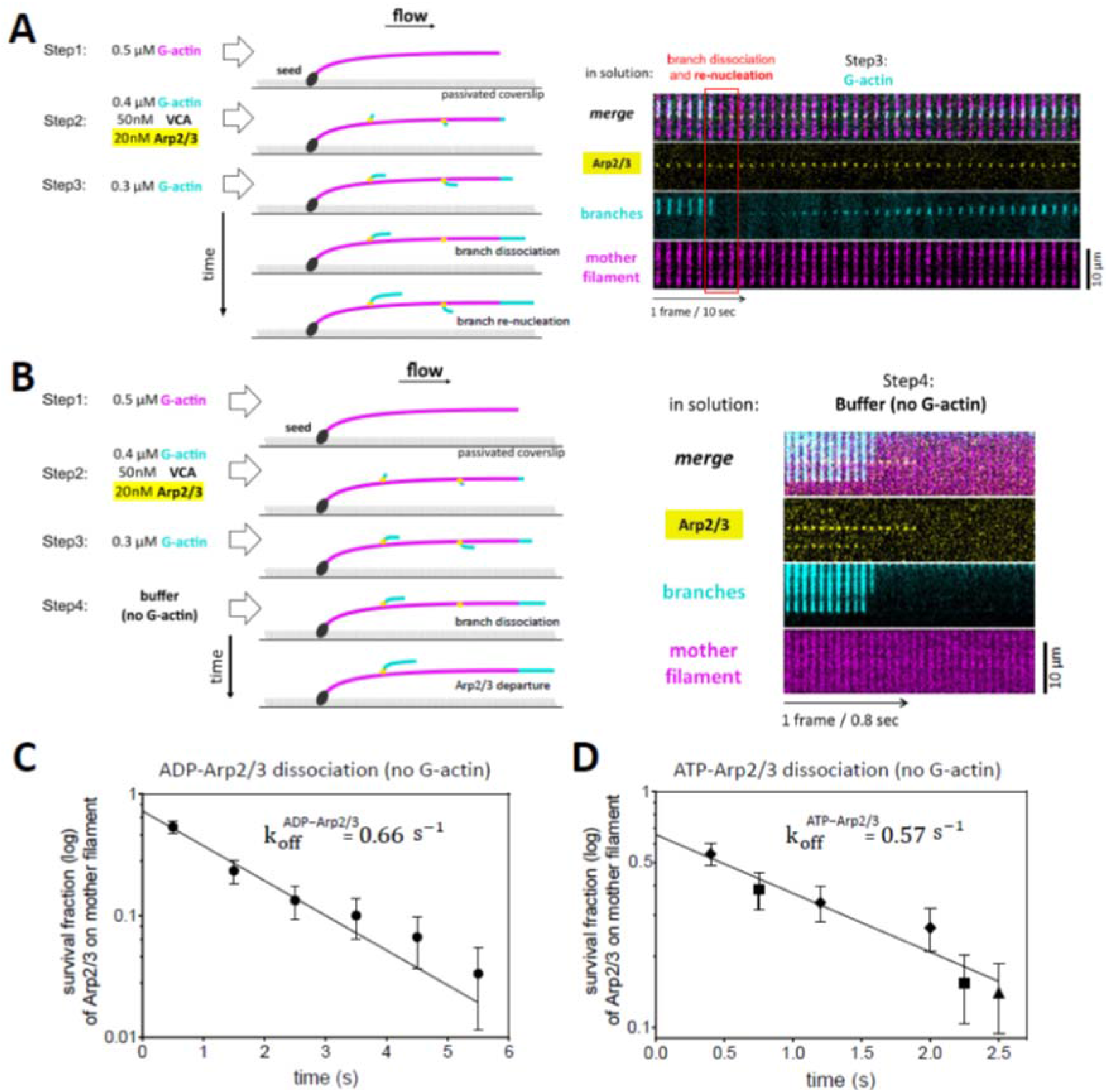
Dissociation of Arp2/3 from the mother filament following debranching. (A) Schematic and timelapse of an experiment carried out with Alexa488-labeled Arp2/3 complex, showing branch dissociation and re-nucleation. Similar to experiments with the same conditions and unlabeled Arp2/3, 70% of the dissociated branches (n=23) were re-nucleated. (B) Schematic and timelapse of an experiment carried out with fluorescently-labeled Arp2/3 complex, in the absence of G-actin during the final step (branch dissociation). Note the shorter time interval between images in this experiment, in order to monitor the departure of Arp2/3 from the mother filament, following the dissociation of the branch. (C) Fraction of Arp2/3 bound to the mother filament (log scale) as a function of time, observed as described in panel (B), using an ATP-free buffer in step 4. The data are pooled from 4 experiments (n=6,15,7,2 branches). (D) Fraction of Arp2/3 bound to the mother filament (log scale) as a function of time, observed as described in panel (B), using our standard buffer with 200 µM ATP in step 4. The data are pooled from different experiments using different time intervals: 0.8 seconds (diamonds, 9 experiments with n=4, 5, 8, 7, 4, 4, 14, 10, 2 branches); 1.5 seconds (squares, 7 experiments with n=14, 14, 4, 3, 7, 1, 9 branches); 5 seconds (triangle, 2 experiments with n=26, 31). In (C,D) time t=0 corresponds to the dissociation of the branch.

We then performed experiments in the absence of G-actin (buffer only) during the debranching step, to monitor the dissociation of the Arp2/3 complex from the mother filament, following the dissociation of the branch (Fig 5B). Experiments were performed in an ATP-free buffer to measure the off-rate of ADP-Arp2/3 (Fig 5C), as well as in our standard buffer (200µM ATP) to measure the off-rate of ATP-Arp2/3 (Fig 5D). We found similar dissociation rates for both situations: k_off_= 0.66 (±0.16) s^-1^ for ADP-Arp2/3 (Fig 5C), and 0.57 (±0.22) s^-1^ for ATP-Arp2/3 (Fig 5D). Knowing the off-rate for ADP-Arp2/3, and considering that the branch re-nucleation ratio is within 5% of the plateau value with 1.2 µM ATP (Fig 4E, based on our error bars) we can determine that the effective rate constant for ATP reloading is greater than 6 µM^-1^s^-1^. This ensures that, with 200 µM ATP, nucleotide exchange is infinitely fast compared to the dissociation of ADP-Arp2/3, and that it is indeed the dissociation of ATP-Arp2/3 that is observed.

In addition, the exponential fits of the survival curves of mother-remaining Arp2/3 complexes (Fig 5C,D) allow us to estimate (y-intercepts) that approximately 70% of Arp2/3 complexes remained on the mother filament upon debranching, in these experiments where we applied a 2.2-2.9 pN pulling force to the branches (in order to accelerate debranching and minimize the photobleaching of Arp2/3 while we acquired images at a rate of 0.2-1.25 Hz, see Methods). Our estimation of 70% mother-remaining Arp2/3 complexes is consistent with the branch re-nucleation ratios we have measured in this force range (Fig 1F).

### Quantitative model for branch dissociation and re-nucleation

In our description of the different pathways leading either to the dissociation of Arp2/3 from the mother filaments or to the re-nucleation of a branch, we hypothesized that the binding of an actin monomer to the ATP-Arp2/3 complex would stabilize its interaction with the mother filament, and promote the nucleation of a new branch. This is consistent with our observation that the branch re-nucleation ratio increases with the G-actin concentration (Fig 1F). In order to quantify this reaction, we measured the branch re-nucleation ratio over a broader range of G-actin concentrations (Fig 6A). We found that it follows a typical saturation curve, with less than 25% of branches re-nucleating below 0.2 µM G-actin, and more than 85% of branches re-nucleating above 1.5 µM. Knowing the rate constant of the competing reaction (dissociation of ATP-Arp2/3 from the mother filament, Fig 5D) we can fit this curve to estimate that G-actin binds to mother-bound ATP-Arp2/3 with rate constant k_on_= 3.4 (±1.24) µM^-1^s^-1^ and a critical concentration C_c_= 0.14 (±0.2) µM. The fit also yields a plateau value of 97(±3)%, which is an estimation of the fraction of debranching events that leave Arp2/3 bound to the mother filament, in the conditions of this experiment where a 1-2 pN force was applied to the branches.

**Fig. 6.**
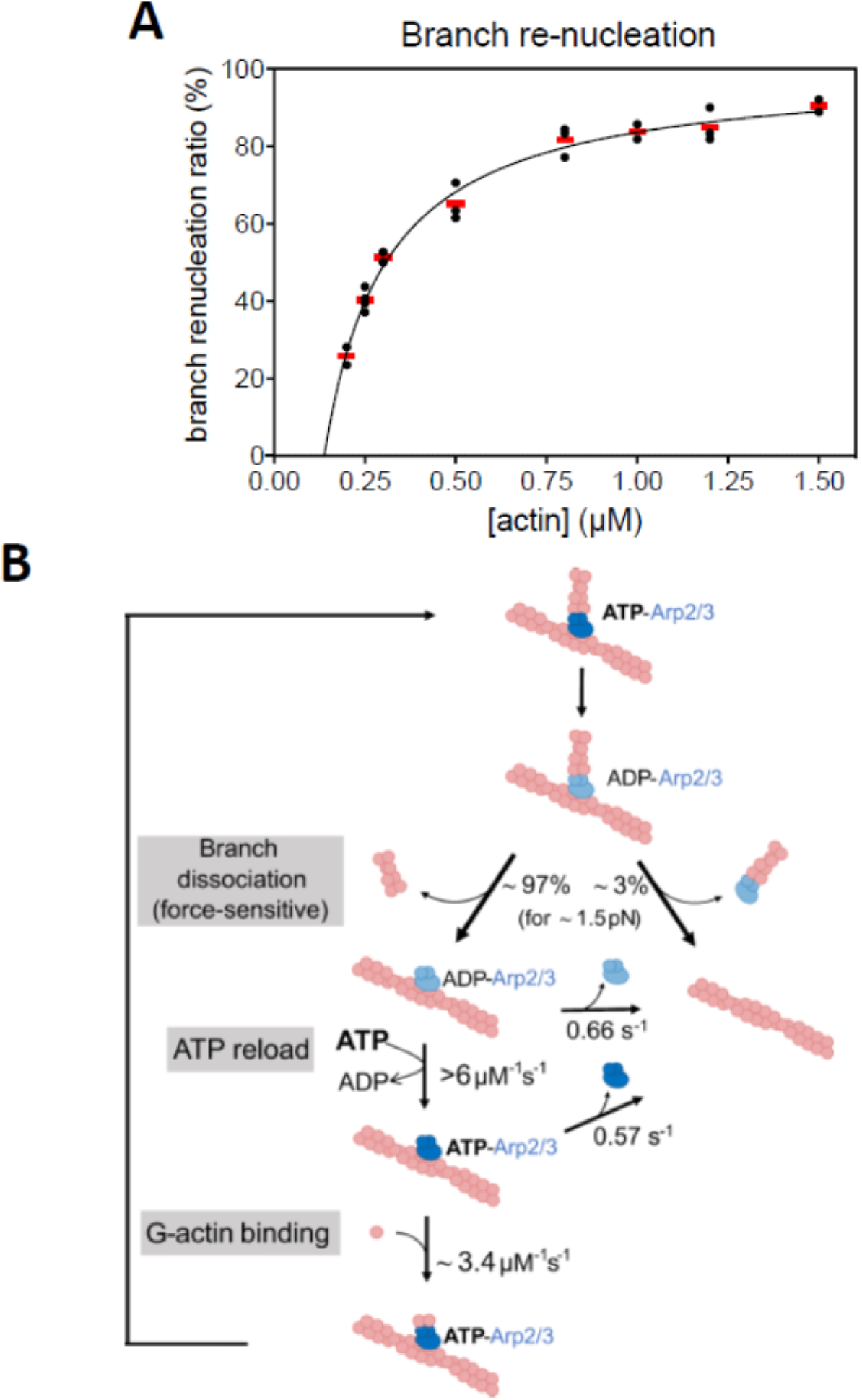
Impact of G-actin concentration on branch re-nucleation, and model for debranching and branch re-nucleation. (**A**) Impact of G-actin concentration on the branch re-nucleation ratio. Each black data point is from a single experiment, monitoring n dissociating branches: n=17, 32 (0.2 µM actin); 35, 32, 38, 32 (0.25 µM); 32, 36 (0.3 µM); 34, 30, 26 (0.5 µM); 32, 36, 35 (0.8 µM); 14, 22 (1 µM); 22, 30, 20 (1.2 µM); 36, 38 (1.5 µM). Red bars indicate average values for each actin concentration. The solid line is a fit, considering that re-nucleation occurs when an actin subunit is added to the Arp2/3 complex before it detaches from the mother filament (see Methods). (**B**) Global model for debranching and branch regrowth. Aged actin filament branches dissociate in a force-sensitive manner, most often leaving an ADP-Arp2/3 complex bound to the mother filament. This mother-remaining Arp2/3 complex will nucleate a new branch, provided that it can load fresh ATP and bind G-actin before detaching from the mother filament.

With these last measurements, we have quantified the final step of the global reaction scheme leading to branch re-nucleation (Fig 6B). This scheme can be summarized as follows. Over time, the Arp2/3 complex in the branch junction hydrolyzes its ATP, and debranching then becomes more likely. This branch dissociation step is accelerated by mechanical forces, which also favor the departure of Arp2/3 from the mother filament. Nonetheless, up to several piconewtons of applied force, the vast majority of debranching events occur with Arp2/3 remaining bound to the mother filament. This mother-remaining ADP-Arp2/3 complex then rapidly exchanges its nucleotide, reloading fresh ATP from solution. If [ATP]>1µM, this reaction always occurs before the dissociation of ADP-Arp2/3 from the mother filament. The mother-bound ATP-Arp2/3 can then bind G-actin from solution, thereby nucleating a new branch that is identical to the initial branch. If the concentration of G-actin is greater than a few micromolars, this reaction always occurs before the dissociation of ATP-Arp2/3 from the mother filament. These rapid reaction rates explain why branches appear to re-nucleate instantly, within the resolution of our experiments (Supp Fig S2).

### Impact of GMF

We wondered whether the global reaction scheme leading to branch re-nucleation (Fig 6B) could be affected by regulatory proteins. To address this question, we monitored branch dissociation and re-nucleation in the presence of Glia Maturation Factor (GMF), a protein known to accelerate debranching (*22*). We used GMF from *Drosophila*, which is very similar to human GMF (Supp Fig S6). We confirmed that GMF accelerates debranching, and we measured that it does so with an apparent dissociation constant K_D_=260 (±117) nM and a maximum debranching rate k_max_=0.071 (±0.008) s^-1^ (for ADP-Arp2/3 and an average pulling force of 1 pN, Fig 7A,B). We also found that GMF drastically hinders branch re-nucleation (Fig 7C). In fact, GMF seemed particularly efficient at preventing branch re-nucleation: in the presence of 25 nM GMF, where we can estimate that [GMF]/([GMF]+K_D_)=9% of branch dissociation events are caused by GMF, the fraction of branches that re-nucleate in the presence of 0.6 µM actin was reduced more than 3-fold (Fig 7C). We verified that GMF did not affect barbed end elongation in our experiments, confirming that GMF does not interact with actin, and indicating that GMF directly binds to Arp2/3 to prevent branch re-nucleation.

**Fig. 7.**
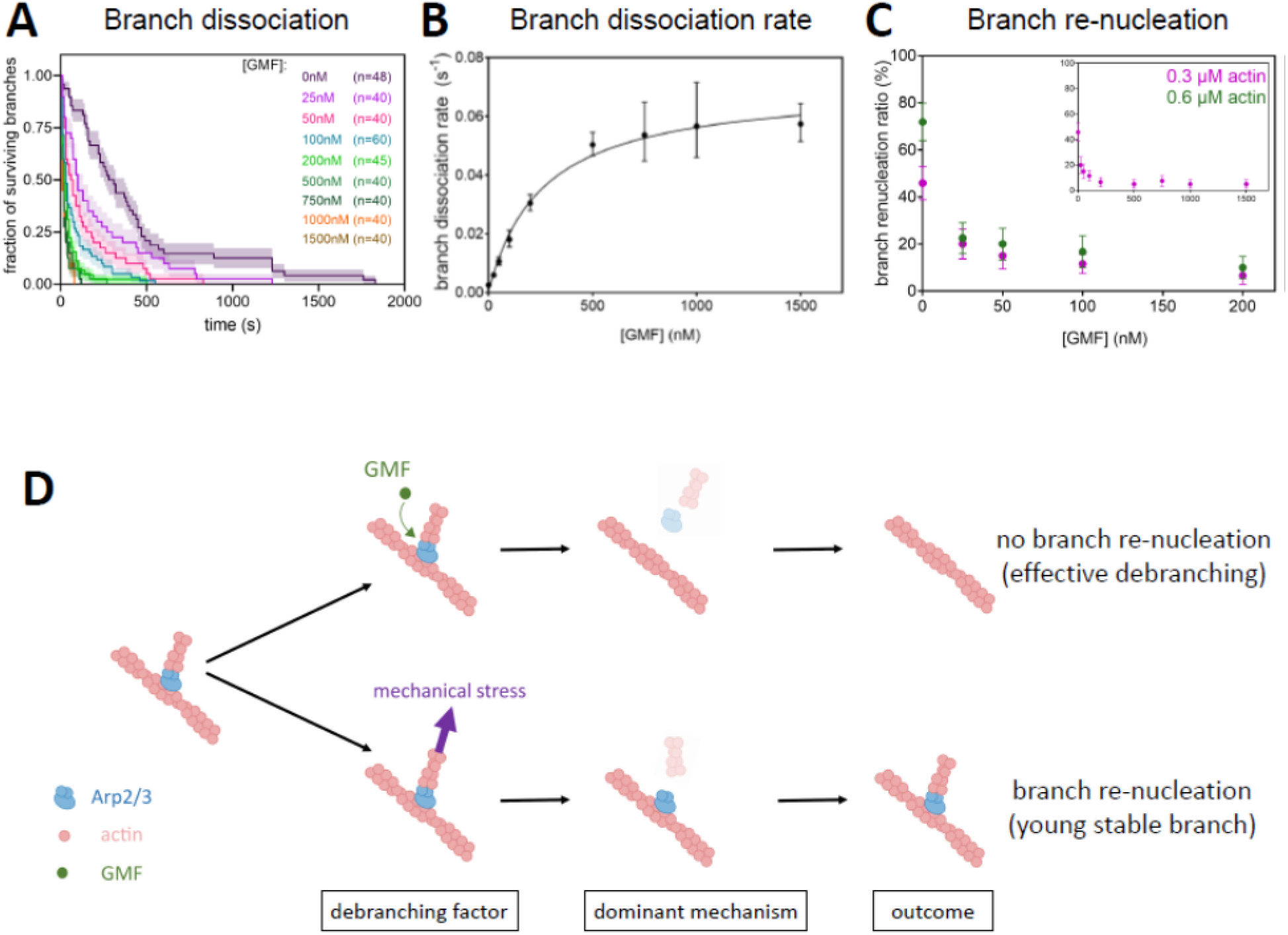
Impact of GMF on branch dissociation and branch re-nucleation. (**A**) Fraction of remaining branches over time, for different GMF concentrations (after 30 minutes of aging, and with an average 1 pN pulling force applied to the branches). (**B**) Branch dissociation rate versus GMF concentration. The rates were determined by exponential fits of the survival curves shown in (A) and the error bars are asymmetrical 95% confidence intervals (Methods). The solid line is a fit by a Michaelis-Menten equation, assuming a rapid equilibrium between GMF and the branch junction. The fit yields a dissociation constant K_D_=260 (±117) nM, a maximum debranching rate k_max_=0.071 (±0.008) s^-1^ and a debranching rate in the absence of GMF comprised between 0 and 0.005 s^-1^ (95% C.I.). (**C**) Branch re-nucleation ratio as a function of GMF concentration, for two different actin concentrations. The data at 0.3 µM actin is from the same data set as panel (A). The data points at 0.6 µM actin each represent an independent experiment, monitoring n=32, 40, 35, 30, 40 dissociating branches, from left to right. Inset: data at 0.3 µM actin shown over a larger range of GMF concentrations. (**D**) Schematic summarizing our interpretation of the different outcomes based on which factor promotes debranching. If branch dissociation is caused by GMF, the dominant outcome is an effective debranching, resulting in a linear filament with no branch. In contrast, if debranching is a consequence of mechanical stress, the dominant outcome is the dissociation of the branch followed by the rapid nucleation of a new branch by the same, ATP-reloaded Arp2/3 complex. This results in a reinforcement of the branch junction.

The impact of GMF on branch re-nucleation contrasts with that of pulling forces, which have a strong effect on the rate of branch dissociation and a moderate impact on branch re-nucleation (Fig 1). It thus appears that the final outcome following a debranching event depends on what factor triggered debranching (Fig 7D). When a branch dissociates because of mechanical stress, the Arp2/3 complex is almost certain to remain bound to the mother filament and nucleate a new branch, and this new branch will be more resistant because it has ATP-Arp2/3 at its junction. In contrast, when branch dissociation is caused by GMF, it is likely that no branch will regrow.

## DISCUSSION

We show here that, in contrast with what is often assumed, the Arp2/3 complex stays bound to the mother actin filament when the branch dissociates. Further, the Arp2/3 that remains on the mother filament can readily nucleate a new branch, from the same location, without requiring reactivation by an NPF. We show that this is a robust mechanism, taking place over a wide range of conditions (Figs 2 and 3). Nonetheless, the likelihood of branch re-nucleation can be reduced by pulling on the branch junction, and by decreasing the concentration of ATP or G-actin. This reflects the three requirements for branch re-nucleation (Fig 6B): first, Arp2/3 must stay on the mother filament upon debranching (slightly less likely when pulling on the branch); second, it must then exchange its nucleotide before detaching (less likely at very low ATP concentration); third, the ATP-Arp2/3 complex must bind actin monomers before detaching (less likely at very low G-actin concentration). In contrast to these factors, which require extreme conditions (high forces, or nearly total depletion of ATP or G-actin) to impact branch re-nucleation, GMF at moderate concentration efficiently prevents the re-nucleation of dissociated branches (Fig 7).

### Molecular insights

Quite strikingly, our data indicate that the dissociation of the branch does not inactivate the Arp2/3 complex remaining on the mother filament. During the initial activation of the Arp2/3 complex, binding to the mother filament can be viewed as a later step, taking place after key conformational changes have been imparted by the binding of the NPF together with G-actin (*10, 28*). This implies that, after the branch has dissociated, the mother-Arp2/3 interactions are able to maintain the Arp2/3 complex in a quasi-active conformation, where it only needs to reload ATP in order to efficiently nucleate a new branch.

The presence of ATP in Arp2/3 is required to nucleate the initial branch (*15, 16*) and ATP hydrolysis promotes branch dissociation, which we show here to result predominantly from the rupture of the Arp2/3-daughter interface. It thus seems that the nucleotide state of Arp2 and Arp3 is key for their interaction with actin. In contrast, the nucleotide state of Arp2 and Arp3 does not seem to directly affect the interaction of the Arp2/3 complex with the mother filament. For instance, after debranching, loading ATP does not stabilize the interaction of the Arp2/3 complex with the mother filament (Fig 5C,D). What seems to stabilize the interaction with the mother filament is the binding of actin to Arp2 and Arp3, which thus appears to trigger an allosteric conformational change of the Arp2/3 complex. At this stage, it is unclear whether binding one actin monomer to Arp2 and Arp3 suffices to stabilize the Arp2/3 complex on the mother filament, or if more actin monomers are required.

In the lone Arp2/3 complex that remains on the mother filament when the branch dissociates, Arp2 and Arp3 exhibit a behavior that is reminiscent of both G-actin and F-actin. The rapid reloading of ATP (Fig 4), similar to what has been reported for free Arp2/3 (*15, 16*), suggests that Arp2 and Arp3 should be in an open G-actin-like conformation, with accessible nucleotide-binding pockets. This seems to change once the Arp2/3 complex is loaded with ATP, as it can bind actin subunits with an on-rate that is only 3-fold lower than that of a filament barbed end (Fig 6). It thus appears that Arp2 and Arp3 are then in a conformation similar to that of a filament barbed end, where the two terminal actin subunits adopt an F-actin-like conformation, as recently shown by cryo-EM (*35*). At this stage, we do not know whether it is the nucleotide state of Arp2 or Arp3 that determines the behavior of the Arp2/3 complex during branch re-nucleation. Early reports suggest that ATP is hydrolyzed more rapidly in Arp2 (*15*) but recent cryo-EM data on yeast Arp2/3 suggest that it could rather be in Arp3 (*29*).

Our quantification of the branch re-nucleation ratio provides decisive insights into the relative strength of the mother-Arp2/3 and Arp2/3-daughter interfaces. Within the range of forces we have studied, Arp2/3 remains bound to the mother filament during the vast majority of debranching events, showing that the mother-Arp2/3 interface is the stronger. The acceleration of debranching with force indicates an overall “slip bond” behavior, where interactions are destabilized by force. The fact that force moderately affects the branch re-nucleation ratio indicates that both interfaces are similarly affected by force, and thus that they both behave like slip bonds. Yet, since the branch re-nucleation ratio decreases with force, it seems that the mother-Arp2/3 interface is slightly more sensitive to force than the Arp2/3-daughter interface.

Our data also show that the Arp2/3-daughter interface is weaker than the interface between actin subunits within the filament. This is consistent with our recent observations on linear (i.e., non-branched) filaments nucleated by Arp2/3 after activation by SPIN90 (*25*). In this situation, the need for a mother filament is bypassed and Arp2/3 is bound simultaneously to SPIN90 and to the pointed end of the nucleated filament (*36, 37*). We observed that the interface between Arp2/3 and the filament ruptured first, leaving Arp2/3 bound to SPIN90 with no actin subunits (*25*). This indicates that the interface between the SPIN90-activated Arp2/3 complex and the nucleated filament, which is very similar to the Arp2/3-daughter interface in branch junctions (*28*), is weaker than the interface between actin subunits. The weakness of the Arp2/3-daughter interface could stem from the specific conformation adopted by the D-loop of the first two actin subunits of the daughter filament in contact with Arp2 and Arp3, as recently observed in branch junctions using cryo-EM (*29*).

Our results showing that GMF prevents branch re-nucleation are reminiscent of our recent observations where GMF enhanced the detachment of Arp2/3 from both SPIN90 and the pointed end of the filament (*25*). They also provide new information on the interaction of GMF with the Arp2/3 complex. Co-crystal structures from GMF and ATP-Arp2/3 show that GMF binds primarily to Arp2 (*23*) and possibly also to Arp3 (*38*), and both of these interactions should destabilize the interface between Arp2/3 and the daughter filament. Our data indicate that GMF is more efficient at preventing branch re-nucleation than at accelerating branch dissociation: the half-maximum effect is reached with less than 25 nM GMF for branch re-nucleation, and with 200 nM GMF for branch dissociation (Fig 7). This suggests that different binding sites could be involved for the two mechanisms, as GMF would interfere with more than one step in our global reaction scheme (Fig 6B).

We also show that, while branch re-nucleation is the dominant outcome for branches made using mammalian Arp2/3 and mammalian actin, it is rare if branches are made using budding yeast Arp2/3 and mammalian actin (Supp Fig S4B). Together with our observation that branches made with yeast Arp2/3 and mammalian actin also dissociate faster (Supp Fig S4A), this suggests that the interface between the mammalian actin mother filament and the yeast Arp2/3 complex is weaker than in an all-mammalian branch junction. Consistently, fission yeast Arp2/3 was recently reported to dissociate from mammalian actin mother filaments upon debranching (*21*). This result possibly reflects a weaker interaction between proteins from evolutionarily distant species. Alternatively, it could reflect a specific difference between branch junctions in yeast and in mammals. Recent cryo-EM data highlight structural differences between fission yeast and mammalian Arp2/3 at branch junctions, but so far they report no major difference regarding the mother-Arp2/3 interface (*28, 29*).

### Physiological implications

In animal cells, the cytoplasm typically contains 100-200 µM G-actin and millimolars of ATP (*34, 39*). These concentrations are far above the ranges where G-actin and ATP are limiting factors for branch re-nucleation (Fig 4D,E and Fig 6A). We thus expect that branch dissociation events would nearly all be followed by branch re-nucleation events, unless they were triggered by a regulatory protein like GMF (Fig 6B). Indeed, the fact that GMF and mechanical forces both accelerate debranching but have a very different impact on branch re-nucleation, lead us to propose that the fate of the Arp2/3 complex largely depends on what factor triggers debranching (Fig 7B).

If debranching is caused by the regulatory protein GMF, it is then very unlikely that a branch will grow back. In this situation, GMF would control the evolution of the branched network towards a more linear architecture, as observed in the rear of lamellipodia or when stress fibers emerge from the cell cortex (*40, 41*). Indeed, a fully effective debranching, where no new branch is re-nucleated, appears essential in order to fully remodel or disassemble the filament network. As such, we propose that the prime role of GMF in cells may be to prevent branches from growing back, redefining its importance for the control of network architecture and disassembly. Whether and how other proteins affecting branch stability, such as coronin or cortactin, would affect branch re-nucleation is now an open question. Other factors should also be investigated, such as the isoform composition of the Arp2/3 complex, which was shown to affect branch stability and its vulnerability to oxidation (*26, 27*).

In addition, our results on the weakness of the Arp2/3-daughter interface in branch junctions indicate that the pointed end of the dissociated branch is unlikely to be capped by Arp2/3. We thus propose that the pointed end of dissociated branches may depolymerize or reanneal with barbed ends, thereby contributing to the reorganization of the network.

In contrast to GMF-induced debranching, if the branch dissociates because of mechanical forces applied to the branch junction, then a new branch is likely to grow back. This is a frequent situation in cells, as branched networks can be exposed to significant mechanical stress. In a lamellipodium, for example, global compression can result in forces with a variety of orientations at the scale of individual branch junctions (*1*). A pushing force exerted on an individual branch, pushing toward the back of the network, would apply a mechanical load on the branch junction similar to the high-angle tension we applied in Fig 2. In a tensed cortex, the combined action of myosin motors and filament crosslinkers could pull on branch junctions in all directions. Since the orientation of the force has a mild impact on the outcome (Fig 2), branch re-nucleation could occur in a broad range of situations.

The forces we have applied in our experiments, in the picoNewton range, are comparable to what can be expected for single filaments in cells. Moreover, since debranching is greatly accelerated by force, an ADP-Arp2/3 branch exposed to an increasing load is likely to dissociate rapidly, when the force is still moderate. It will then leave behind an Arp2/3 complex that will rapidly be “rejuvenated” by ATP and from which a new branch will grow. The network would thus rapidly shed old, fragile branches and replace them with new, more resistant branches. This is a new mechanism by which branched actin networks could adapt to mechanical stress by self-repairing, reorganizing and strengthening. Importantly, branch re-nucleation events can take place away from NPF-decorated surfaces. In the context of the lamellipodium, this could explain why growing barbed ends can sometimes be observed away from the leading edge (*42*–*44*).

Branched actin networks are found in a variety of cellular contexts, with different turnover rates and different functional requirements. Future studies should uncover how the rejuvenation mechanism endowed by branch re-nucleation contributes to their maintenance and turnover.

## Acknowledgments

We thank Alphée Michelot for the gift of Arp2/3 purified from budding yeast, Pekka Lappalainen for the gift of GMF plasmid, and Mohan Balasubramanian for the gift of the *P. pastoris* strain. We thank Tamara Advedissian, Arnaud Echard, Cécile Leduc and Hugo Wioland for reading an earlier version of this manuscript and making useful suggestions.

## Funding

European Union’s Horizon 2020 Marie Sklodowka-Curie individual fellowship program H2020-MSCA-IF-101028239-MolecularArp (LC).

Cancer Research UK CC2096 core funding to the Francis Crick Institute (MM, MW)

UK Medical Research Council CC2096 core funding to the Francis Crick Institute (MM, MW)

Wellcome Trust CC2096 core funding to the Francis Crick Institute (MM, MW)

European Research Council (ERC) under the European Union’s Horizon 2020 research and innovation programme, grant agreement No 810207 (MW)

European Research Council (ERC) grant StG-679116 (AJ)

Agence Nationale de la Recherche grant RedoxActin (GRL)

Fondation pour la Recherche Medicale grant EQU202203014630 (GRL)

## Author contributions

Conceptualization: AJ, GRL

Resources: LC, MM, BG

Methodology: FG, LC, MW, AJ, GRL

Formal analysis: FG, AJ, GRL

Investigation: FG

Visualization: FG, AJ, GRL

Funding acquisition: LC, MW, AJ, GRL

Project administration: AJ, GRL

Supervision: AJ, GRL

Writing – original draft: AJ, GRL

Writing – review & editing: FG, LC, MW, AJ, GRL

## Competing interests

Authors declare that they have no competing interests.

## Supplementary Materials

Materials and Methods

Figs. S1 to S6

References 45-51

## Notes

### Competing Interest Statement

The authors have declared no competing interest.

