## Supplementary material for "Regeneration of actin filament branches from the same Arp2/3 complex": methods and supp figs

**The PDF file includes:**

Materials and Methods

Figs. S1 to S6

**Materials and Methods**

**Buffers and proteins**

**Proteins**

Skeletal muscle actin (Uniprot P68135) was purified from rabbit muscle acetone powder following the protocol from (*45*) as described in (*33*).

Recombinant cytoplasmic Gamma-actin (Uniprot P63261) was expressed in *Pichia pastoris*, with N-terminal acetylation and His73 methylation, following the protocol described in (*46*). The *P. pastoris* strain, expressing both NAA80 acetyltransferase and SETD3 methyltransferase, was a kind gift from the Balasubramaniam lab.

Spectrin-actin seeds from human red blood cells were purified as described in (*33*).

Arp2/3 complex was purified from sheep brain, as previously described in (*47*). For assays using fluorescently labeled Arp2/3 (Figure 5), we used recombinant Arp2/3 produced in Sf9 insect cells, produced and purified as detailed in (*24*). Arp2/3 from *Saccharomyces cerevisiae* was a kind gift from Alphée Michelot.

Recombinant human profilin1 (Uniprot P07737) was expressed and purified as described in (*48*).

Recombinant N-terminal GST tag human N-WASP-VCA (aa 392–505, Uniprot O00401) and human WASP-VCA (418–502, Uniprot P42768) were expressed and purified as described in (*25*).

Recombinant C-terminal His-tagged drosophila GMF (full length, aa 1–138, Uniprot Q9VJL6, a kind gift from the Lappalainen lab) was expressed and purified as described in (*25*).

**Protein labeling**

Actin was fluorescently labeled on the surface lysine 328, using Alexa Fluor 488, Alexa Fluor 568, or Alexa Fluor 647 NHS ester (Thermo Fisher), or ATTO 643 NHS ester (Atto-tec) as described in detail in (*49*).

Arp2/3 was fluorescently labeled using Alexa Fluor 488 or Alexa Fluor 568 C_5_-maleimide (Thermo Fisher). The protein solution was prepared for labeling by performing a buffer exchange to remove DTT from the solution. This was accomplished by passing the protein solution through a MicroBiospin 6 column (BioRad). The exchange buffer contained 20 mM HEPES at pH 7.2, 0.2 mM MgCl2, and 0.2 mM ATP. The buffer exchange was performed by centrifuging the sample at 1000 g for 4 minutes. Next, a 10-fold excess of Alexa 488 (or Alexa 568) C_5_-maleimide dissolved in DMSO was added to the Arp2/3 solution and incubated on ice for 1 hour. The reaction was stopped by adding 1mM DTT to the solution. To remove any unreacted excess dye, a MicroBiospin 6 column (BioRad) was used by centrifugation at 1000 g for 4 minutes. We obtained on average 3.5 Alexa dyes per Arp2/3 complex.

**Buffer**

All microfluidics experiments were performed in F-buffer containing: 5 mM Tris HCl pH 7.0, 50 mM KCl, 1 mM MgCl2, 0.2 mM EGTA, 0.2 mM ATP, 10 mM DTT, 1 mM DABCO, supplemented with 0.1% Bovine Serum Albumin (BSA).

**Experiments**

**Data Acquisition**

Observations were made using a Nikon TiE inverted microscope equipped with a 60x oil-immersion objective and an Evolve 512 EMCCD camera (Photometrics). The temperature of the microfluidics chamber was maintained at 25 °C (± 0.2 °C) using a collar objective heater (Okolab). We used an azimuthal total internal reflection fluorescence (TIRF) illumination setup (iLAS2, Gataca-system), with 488, 561 ad 642 nm tunable lasers (max. power 100 mW each). The manipulation and adjustment of the TIRF setup were carried out using Metamorph software (Version 7.10.4.407). Image analysis was performed using Fiji software (*50*).

**Basic microfluidics experiment (Fig 1)**

Microfluidics experiments were conducted using Poly-Dimethyl-Siloxane (PDMS, Sylgard) chambers based on the original protocol from (*30*), described in detail in (*51*).

Spectrin-actin seeds were flowed in the microfluidics chamber at 20 pM for 2 minutes. Next, the surface was passivated by exposing it to a solution containing 5% BSA (bovine serum albumin) for 10 minutes. After passivation, surface-anchored mother filaments were polymerized by flowing in 0.6 μM 10% Alexa-488 labeled G-actin for 5 to 10 minutes. Next, to induce the nucleation of actin filament branches at a sufficient density, mother filaments were exposed to a solution of 20 nM Arp2/3, 50 nM VCA, and 0.4 μM 10% Alexa-568 labeled G-actin for 45 seconds to 2 minutes. However, it should be noted that, in all experiments, the indicated time of exposure to the branching solution refers to the period following the increase of the flow rate, after the reservoir containing the protein solution is plugged to the microfluidic device. During this time, mother filaments were exposed to a protein concentration that progressively increased from zero to 70% (after 45 seconds) to 100% (after 2 minutes) of the nominal concentration (*51*). Afterwards, actin filament branches and mother filaments were aged with a low flow of 0.3 or 1 μM 10% Alexa-568 labeled G-actin for 20 minutes. And finally, they were exposed to debranching conditions: 0.3 μM 10% Alexa-568 labeled G-actin with different flow rates (to apply different forces). The dissociation of branch filaments and their renucleation were observed over time and acquired with the acquisition rate of 1 frame every 10 seconds.

**Experiment on force orientation (Fig 2)**

Spectrin-actin seeds were flowed to the microfluidics chamber at 20 pM for 2 minutes. Then the surface was rinsed by buffer and passivated by 0.5% BSA and 0.1% biotin-BSA for 10 to 15 minutes. Surface-anchored mother filaments were polymerized in three steps sequentially, to create three segments: starting from the seed, first a non-biotin-actin segment (by 0.8 μM 10% Alexa-488 labeled G-actin), then a biotin-actin segment (by 0.8 μM 6% Alexa-488 labeled G-actin, 50% to 75% biotin labeled), and finally a short non-biotin actin segment (by 0.8 μM 10% Alexa-488 labeled G-actin). Next, the filaments were exposed to 20 to 50 nM neutravidin for 2 minutes. Afterwards, mother filaments were exposed to 20 nM Arp2/3, 50 nM VCA, and 0.4 μM 10% Alexa-568 labeled G-actin for 45 seconds to 2 minutes to nucleate actin filament branches at sufficient density. Then the actin filament branches, as well as mother filaments, were elongated with a low flow of 0.3 μM 10% Alexa-568 labeled G-actin for 4 minutes. Finally, they were exposed to debranching conditions: 0.3 μM 10% Alexa-568 labeled G-actin, flowing perpendicular to the mother filament orientation (from left-hand side channel to all three other channels). The dissociation of branch filaments and their re-nucleation were recorded over time at 1 frame every 10 seconds. For each branch, the angle between the branch (local direction of the flow) and the mother filament at the branch junction was measured manually using the Fiji software.

**Experiment on unlabeled cytoplasmic gamma-actin (Fig 3C)**

Spectrin-actin seeds were flowed into the microfluidics chamber at 20 pM for 2 minutes. Then the surface was rinsed with F-buffer and passivated by being exposed to a solution containing 5% BSA, for 10 minutes. After rinsing, surface-anchored mother filaments were polymerized from spectrin-actin seeds in three steps sequentially, to create three segments: starting from the seed, first a labeled alpha-actin segment (by 0.6 μM 10% Alexa-488 labeled alpha G-actin), then an unlabeled gamma-actin segment (by 0.6 μM unlabeled gamma G-actin), and then again a labeled alpha-actin segment (by 0.6 μM 10% Alexa-488 labeled alpha G-actin). Next, the mother filaments were exposed to 20 nM Arp2/3, 50 nM VCA, and 0.4 μM unlabeled gamma G-actin for 1 minute. Afterwards, the filaments and branched junctions were aged while exposed to a solution of 0.3 μM 10% Alexa-568 labeled alpha G-actin, circulating at a low flow rate for 20 minutes. Finally, they were exposed 0.3 μM 10% Alexa-568 labeled G-actin, at a flow rate that applied a 1-1.5 pN force to the branch junction. For comparison, we monitored over time (acquisition rate of 1 frame every 10 seconds) the debranching and re-nucleation of branches that were nucleated from the side of mother filaments, either along labeled alpha-actin or unlabeled gamma-actin segments.

**Experiment on profilin-actin (Fig 3D)**

Spectrin-actin seeds were flowed into the microfluidics chamber at 20 pM for 2 minutes. Then the surface was rinsed with F-buffer and passivated by being exposed to a solution containing 5% BSA for 10 minutes. After rinsing, surface-anchored mother filaments were polymerized by 0.6 μM 10% Alexa-488 labeled G-actin. Next, mother filaments were exposed to 20 nM Arp2/3, 50 nM VCA, and 0.4 μM 10% Alexa-568 labeled G-actin for 1 minute to nucleate actin filament branches.In the next step, two experimental solutions (actin alone or equimolar amounts of actin and profilin) were introduced into the microfluidics chamber, side by side at equal flow rates, resulting in each solution occupying half of the chamber. By alternating stage positions during the acquisitions of images, this experimental setup allowed us to simultaneously investigate two distinct conditions within the same chamber while keeping all other variables constant. The dissociation of branch filaments and their re-nucleation, for each debranching condition, were observed over time and acquired with the acquisition rate of 1 frame every 10 seconds.

**Experiment on comparison of the first and second generation (Fig 4A, B)**

The spectrin-actin seeds were introduced into the microfluidics chamber at a concentration of 20 pM and allowed to flow for 2 minutes. Subsequently, the surface was rinsed with F-buffer and passivated by being exposed to a solution containing 5% BSA for a duration of 10 minutes. After rinsing, surface-anchored mother filaments were polymerized by 0.8 μM 10% Alexa-488 labeled G-actin. Next, mother filaments were exposed to 20 nM Arp2/3, 50 nM VCA, and 0.4 μM 10% Alexa-568 labeled G-actin for 45 seconds to nucleate actin filament branches. Then branches and mother filament were exposed to 0.4 μM 10% Alexa-568 labeled G-actin for 90 seconds to elongate nucleated branches. Afterwards, branches were aged for 20 minutes with a low flow of 0.18 μM 10% Alexa-568 labeled G-actin. Finally, branches were exposed to debranching conditions: 0.3 μM 10% Alexa-568 labeled G-actin. The dissociation of branch filaments, renucleation of second-generation branches, and their dissociation were observed over time and acquired with the acquisition rate of 1 frame every 10 seconds.

**Experiment on ATP depletion by resin treatment (Fig 4D)**

ATP in the solution of actin in FME buffer was removed by two consecutive treatments (2x15 minutes) with 5% (w/w) Ion Exchange Resins (AG 1-X8 Resin, Bio-Rad).

Spectrin-actin seeds were flowed to the microfluidics chamber at 20 pM for 2 minutes. Then the surface was rinsed with F-buffer and passivated by exposing it to a solution containing 5% BSA for 10 minutes. After rinsing, surface-anchored mother filaments were polymerized by 0.7 μM 10% Alexa-488 labeled G-actin. Then mother filaments were exposed to 20 nM Arp2/3, 50 nM VCA, and 0.4 μM 10% Alexa-568 labeled G-actin for 1 minute to nucleate actin filament branches. Next, two experimental solutions were flowed side by side to the microfluidics chamber at equal flow rates so that each of them only occupied half of the chamber. Experimental solutions were resin-treated solution (1 μM 10% Alexa-568 labeled G-actin, [ATP] < 1 μM), rescued resin-treated solution of actin (1 μM 10% Alexa-568 labeled G-actin , ATP added after treatment), and control (1 μM 10% Alexa-568 labeled G-actin, [ATP] = 200 μM). The dissociation of branch filaments and their re-nucleation, for each debranching condition, were observed over time and acquired with the acquisition rate of 1 frame every 10 seconds.

**Experiment on labeled Arp2/3 (Fig 5)**

The chamber was incubated with a solution of 20 mg/ml PLL-PEG in PBS for 90 minutes. Next, it was incubated with 100 pM Spectrin-actin seeds for 2 minutes. To enhance passivation, the chamber was further incubated with a solution containing 1% BSA and 0.5 mg/ml casein for 30 more minutes. After rinsing, surface-anchored mother filaments were polymerized by 0.7 μM 10% Alexa-647 labeled G-actin. Then, mother filaments were exposed to 20 nM Alexa-568 (or 488) labeled Arp2/3, 50 nM VCA, and 0.4 μM 10% Alexa-488 (or -568) labeled G-actin for 45 seconds to nucleate actin filament branches. Filaments were aged with 0.3 μM 10% Alexa-488 (or -568) labeled G-actin for 30 minutes. Finally, filaments were exposed to a high flow of FME buffer containing either 200 μM ATP or 200 μM ADP. The dissociation of branch filaments and departure of labeled Arp2/3 molecules were observed over time (at 50% laser power) and recorded with an acquisition rate of 1 frame every 0.8, 1.5, or 5 seconds in the presence of ATP, or 1 frame per second in the presence of ADP.

**Experiment on actin concentration vs re-nucleation ratio (Fig 6A)**

Spectrin-actin seeds were flowed into the microfluidics chamber at 20 pM for 2 minutes. The surface was then rinsed with buffer and passivated for 10 minutes using 5% BSA. Following this, mother filaments anchored to the surface were polymerized using 0.8 μM 10% Alexa-488 labeled G-actin. Mother filaments were exposed to 20 nM Arp2/3, 50 nM VCA, and 0.4 μM 10% Alexa-568 labeled G-actin for 45 seconds to 1 minute in order to nucleate actin filament branches. Next, mother filament and branches were elongated and aged with a low concentration of actin (0.3 or 0.4 μM 10% Alexa-568 labeled G-actin) for 30 minutes. And finally, they were exposed to debranching conditions: different concentrations of 10% Alexa-568 labeled G-actin. The average force for each experiment was calculated by determining the average length of branches and the total flow rate in the chamber. Flow rates were adjusted to maintain an average force of 1 to 2 pN. The dissociation of branches and their subsequent renucleation were observed over time and recorded at an acquisition rate of 1 frame per 10 seconds.

To control the length of actin branches during the experiments performed with high actin concentrations (> 1.2 µM), filament and branches were exposed to 1.5 µM of CP for 60 seconds and allowed to age further up to 30 minutes while remaining capped (using 0.3 μM 10% Alexa-568 labeled G-actin), before being exposed to the debranching conditions mentioned above.

**Experiment on the impact of GMF on debranching and re-nucleation (Fig 7)**

Spectrin-actin seeds were flowed into the microfluidics chamber at 20 pM for 2 minutes. Then the surface was rinsed by buffer and passivated by 5% BSA for 10 minutes. After rinsing, surface-anchored mother filaments were polymerized by 0.8 μM 10% Alexa-488 labeled G-actin. Mother filaments were exposed to 20 nM Arp2/3, 50 nM VCA, and 0.4 μM 10% Alexa-568 labeled G-actin for 1 minute to nucleate actin filament branches. Next, mother filament and branches were aged with 0.4 μM 10% Alexa-568 labeled G-actin for 30 minutes. Finally, the chamber was exposed to the debranching conditions: different concentrations of GMF (up to 1500 nM) and 0.3 (and 0.6) μM 10% Alexa-568 labeled G-actin. The dissociation of branch filaments and renucleation of second-generation branches were observed over time and acquired with the acquisition rate of 1 frame every 10 seconds.

**Experiment with a bright fluorescent pointed end region of branches (Fig S3B-D)**

Spectrin-actin seeds were flowed into the microfluidics chamber at a concentration of 20 pM for 2 minutes. The chamber surface was then rinsed with buffer and passivated with 5% BSA for 10 minutes. Before performing the experiment, we performed a speckle experiment.

*Speckle experiment*

In order to determine the fluorescence intensity of a single Alexa488 fluorophore on actin, we monitored filaments sparsely labeled with Alexa488. Mother filaments were polymerized using 0.8 μM labeled G-actin (containing 10% Alexa-568-actin and 0.1% Alexa-488-actin). Then, by testing different laser powers and acquisition times, we determined a set of illumination conditions (100% laser power, 1000 ms acquisition time) that ensure the clear detection of a single Alexa-488 fluorophore in our setup.

*Main experiment*

Starting from a fresh coverslip surface, mother filaments were polymerized using 0.6 μM of 10% ATTO-643 labeled G-actin from spectrin-actin seeds. To nucleate actin filament branches, the mother filaments were exposed to 20 nM Arp2/3, 50 nM VCA, and 0.4 μM of 46.8% Alexa-488 labeled G-actin for 45 seconds. Highly labeled branch junctions were observed with an acquisition rate of 1 frame every 5 seconds. The filaments and branch junctions were aged for 20 minutes in the presence of 0.2 μM of 10% Alexa-568 labeled G-actin, followed by exposure to debranching conditions using 0.5 μM of 10% Alexa-568 labeled G-actin for 10 minutes. Debranching and re-nucleation events were observed and acquired with an acquisition rate of 1 frame every 20 seconds. During that time, the Alexa488 fluorophores were not excited, in order to avoid photobleaching (the 488 nm laser was turned off). Finally, to assess the presence of Alexa-488-actin subunits from the first generation of branches at the junction of re-nucleated branches, filaments were monitored at an acquisition rate of 1 frame every 5 seconds, using all three different wavelengths (488 nm, 561 nm, and 642 nm excitation lasers) and using the illumination conditions determined in the speckle experiment described above to ensure that single Alexa-488 molecules would be detected.

**Data analysis**

**Branch survival functions**

The fraction of surviving branches as a function of time is computed by the Kaplan-Meier method using GraphPad Prism version 9 for Windows. The error bars represent standard error calculated by the method of Greenwood.

**Branch re-nucleation ratio**

The branch re-nucleation ratio represents the ratio of the number of re-nucleated branches to the number of dissociated branches. Error bars show the binomial standard deviation.

**Calculation of the force applied to the branch junction**

The tensile force exerted on branches is a result of the friction (viscous drag) of the fluid flow applied to the filaments. As characterized previously in (*31*), the pulling force on the branch junction is the product of the length of the branch, the local flow velocity, and the longitudinal friction coefficient per unit length of the actin filament (η_actin_ = 6. 10^-4^ pN.µm^-2^.s). In each experiment, the average force and standard deviation of the force were determined for the population of analyzed branches.

Off-rate of Arp2/3 in the presence of ADP or ATP (Fig 5C, D)

The off-rate of lone Arp2/3 (in the presence of ADP or ATP) was derived from the exponential fit of the survival fraction of Arp2/3 on mother filament over time, with a 95% confidence interval, using GraphPad Prism version 9 for Windows.

**On-rate of actin monomers (Fig 6)**

The dependence of the branch re-nucleation ratio as a function of actin concentration was fitted considering the scheme shown in Fig 6B, where branches are re-nucleated when G-actin is added to the ATP-Arp2/3 complex before it detaches from the mother filament:

$$branch renucleation ratio=\frac{A\times k_{on}(C-C_{C})}{k_{on}(C-C_{C})+k_{off}^{ATP-Arp2/3}}$$

where $k_{off}^{ATP-Arp2/3}$is the detachment rate of ATP-Arp2/3 from the side of the mother filament, A is the maximum branch re-nucleation ratio (which also corresponds to the percentage of dissociated branches whose Arp2/3 has remained on the mother filament during branch detachment, and then loaded ATP), $k_{on}$ is the on-rate constant for G-actin binding to the ATP-Arp2/3 complex on the mother filament, and $C_{C}$ is the actin critical concentration for that reaction. A, $C_{C}$ and $k_{on}$ are free parameters during the fitting procedure.

**GMF-induced branch dissociation rates (Fig 7)**

The branch dissociation rate in the presence of GMF was derived from the exponential fit of survival fractions (Fig 7A), with a 95% confidence interval, using GraphPad Prism.

The dependence of the branch dissociation rates as a function of GMF concentration was fitted by the classical Michaelis–Menten kinetics equation, with k (dissociation rate without GMF), k_max_ (maximum dissociation rate), and K_D_ (dissociation constant of GMF on the branch junction) as free parameters:

$$branch dissociation rate=k+\frac{[GMF]}{[GMF]+K_{D}}(k_{max}-k)$$

**Fluorescent intensity of re-nucleated branch at the junction over time (Fig S2)**

The intensity of each individual branch junction was measured as the average intensity in a 4x4-pixel region over time at each time interval. The background intensity in the vicinity of the same branch junction was subtracted from the measured intensity. For each branch, the intensity reaches a plateau when the branch grows out of the 4x4-pixel region, and this plateau is used to normalize the fluorescent signal. The plots in Fig S2B show, for each experiment, the normalized intensity over time, averaged over the observed population of re-nucleated branches.

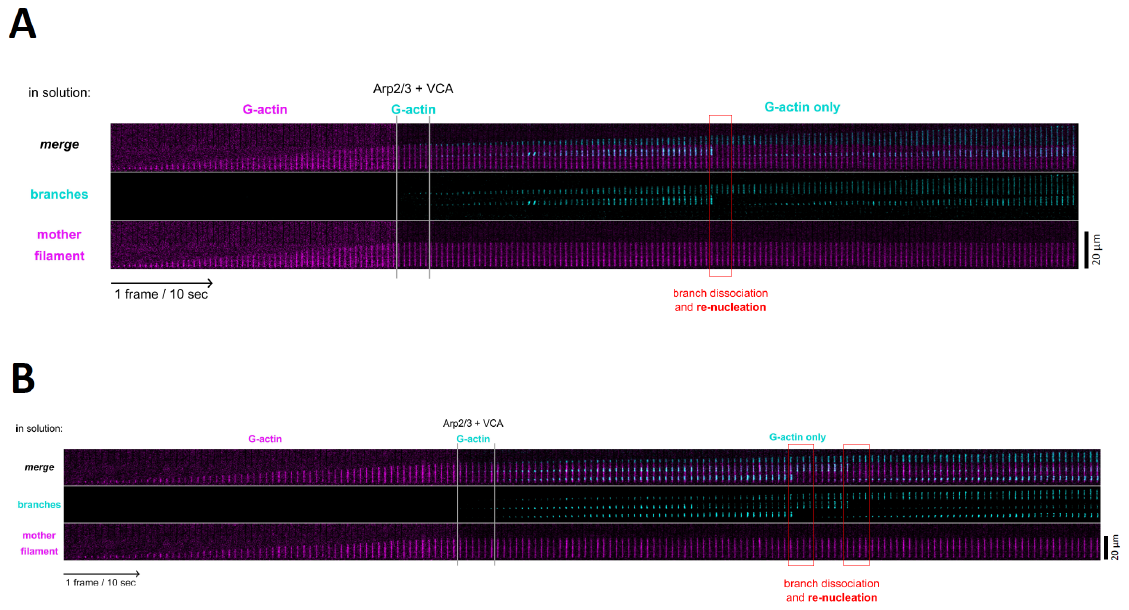

Fig. S1. Additional examples of timelapses showing branch dissociation and re-nucleation, similar to Fig 1C,E, over longer time scales. (A) One branch is visible. It dissociates and another one is re-nucleated at the same location. (B) Two branches are visible on the same mother filament. Each dissociates and each is replaced by a re-nucleated branch. In (A) and (B), the elongation of the barbed end is clearly visible, during the different steps of the experiment. The red boxes indicate branch dissociation and re-nucleation events.

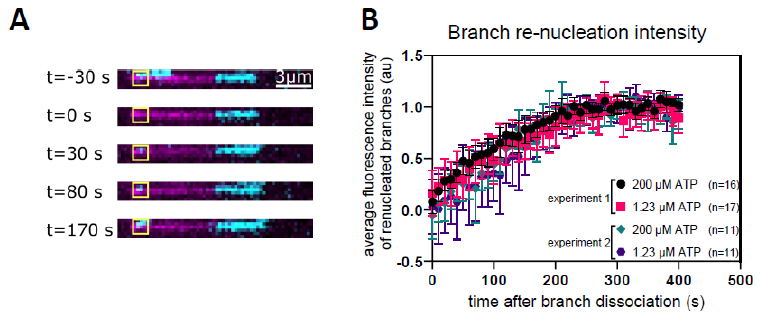

Fig. S2. Branch re-nucleation is observed immediately following branch dissociation.

(**A**) Time-lapse of branch dissociation followed by branch re-nucleation. The average intensity of the branch junction was measured within a 4x4-pixel region, over time. Mother filaments and branches were polymerized with actin labeled by Alexa-488 and Alexa-568, respectively. (**B**) Average fluorescence intensity of n re-nucleated branches over time. Time zero marks branch dissociation, determined as the frame when the branch (Alexa-568) signal is lost. No lag was detected between branch dissociation and branch re-nucleation.

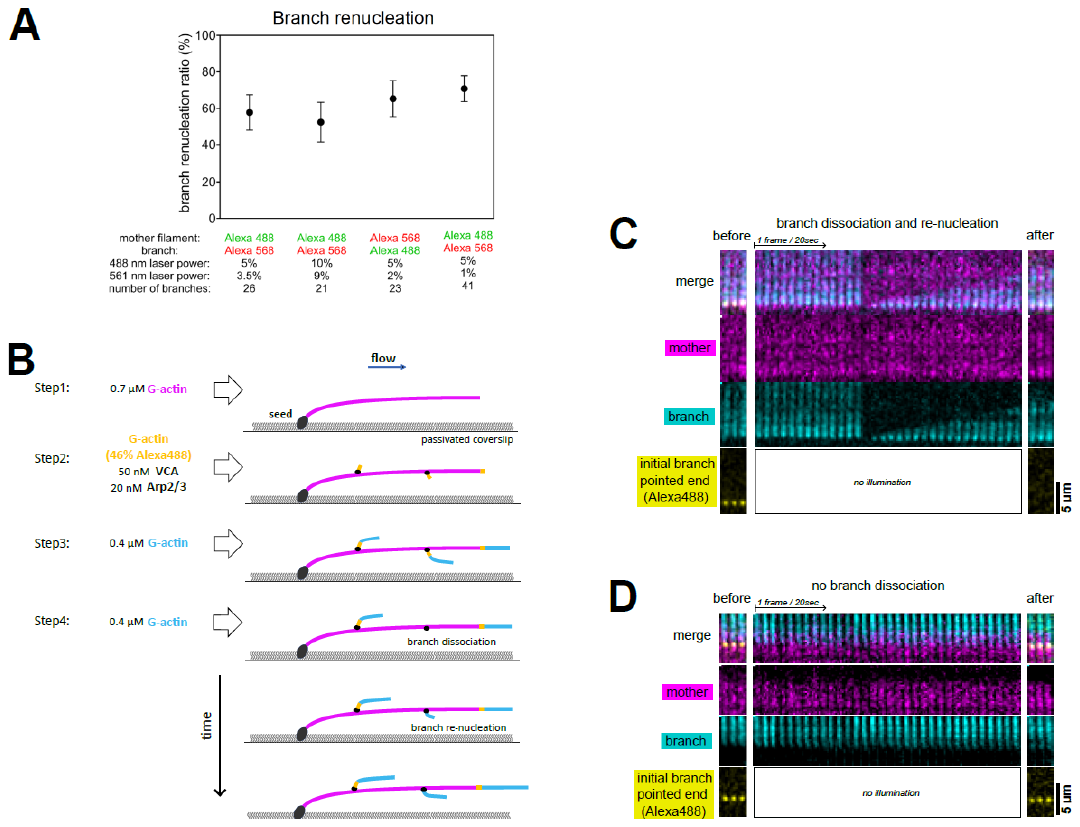

Fig. S3. Branch dissociation events are not due to severing of the branch close to the branch junction. (A) Effect of laser power and labeling on branch re-nucleation. (B) Schematic of experiment designed to look at bright Alexa-488-labeled segments of branches close to the branch junction. Branches with a clearly visible Alexa-488 signal at their branch junction after the nucleation of the initial branch (step 3) were further monitored over time and analyzed (step 4). (C) Time lapse of a branch that dissociated and re-nucleated (during step 4, as shown in (B)), shown as an example: after re-nucleation, the Alexa-488 signal is no longer detectable, in conditions where a single fluorophore would be detected. Nearly none of these re-nucleated branches (1 out of 16) had a detectable Alexa-488 signal at their branch junction at the end of the experiment. (D) Time lapse of a branch that did not dissociate (during step 4, as shown in (B)), shown as an example: the Alexa-488 signal is still visible at the end of the experiment. These branches provide an internal control. Nearly all of them (11 out of 12) still had a detectable Alexa-488 signal at the end of the experiment.

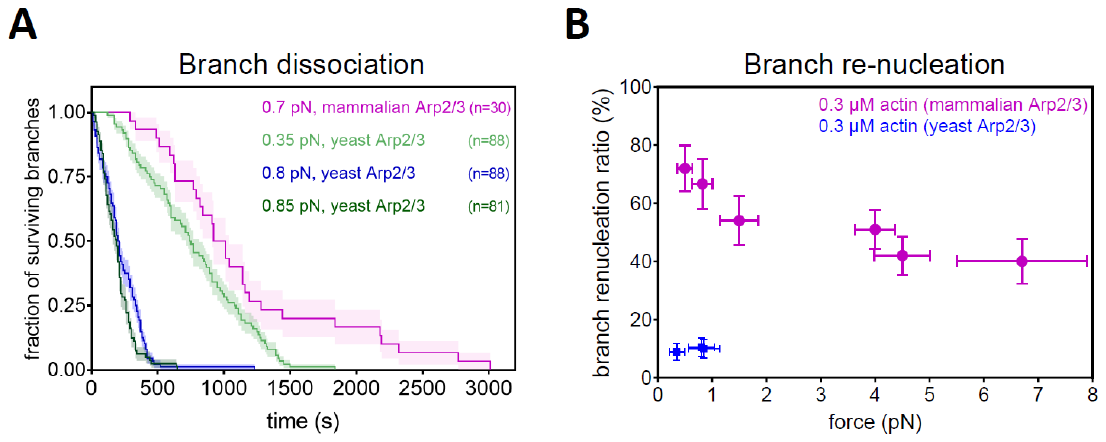

**Fig. S4. Branches nucleated by Arp2/3 from S. Cerevisiae dissociate faster and barely re-nucleate.** (**A**) Comparison of the dissociation of branches nucleated from mammalian Arp2/3 (pink curve) and *S. Cerevisiae* Arp2/3 (other curves), exposed to different pulling forces. Each curve is from a single experiment, where the branches were pre-aged for 4 minutes (before t=0). In all experiments, mammalian alpha-skeletal actin was used. (**B**) Comparison of the ratio of re-nucleated branches, from mammalian Arp2/3 (pink) and *S. Cerevisiae* Arp2/3 (blue), for different pulling forces. The data for mammalian Arp2/3 is the same as in Fig 1F. For yeast Arp2/3, each point is from a single experiment, monitoring, from left to right, n=90, 95, 95 branches.  In all experiments, mammalian alpha-skeletal actin was used.

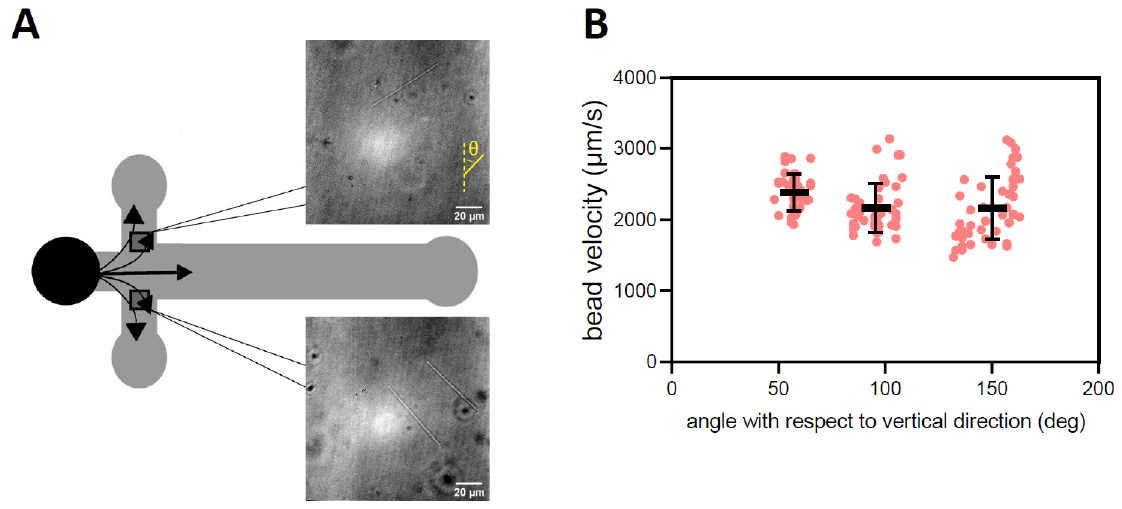

**Fig. S5. Flow velocity as a function of flow direction.** (**A**) The microscope images (transmitted light) show the traces of one bead (top) and 2 beads (bottom) passing in two different regions of the microchamber. The images were acquired over 30 ms and beads have the diameter of 3 μm. (**B**). Each data point indicates the velocity of a single bead, traveling in a direction making an angle θ with respect to the vertical direction on the images. The black marks indicate the average velocities and standard deviations for 3 subpopulations of n=39, 43, 51 beads (from left to right) with similar angles.

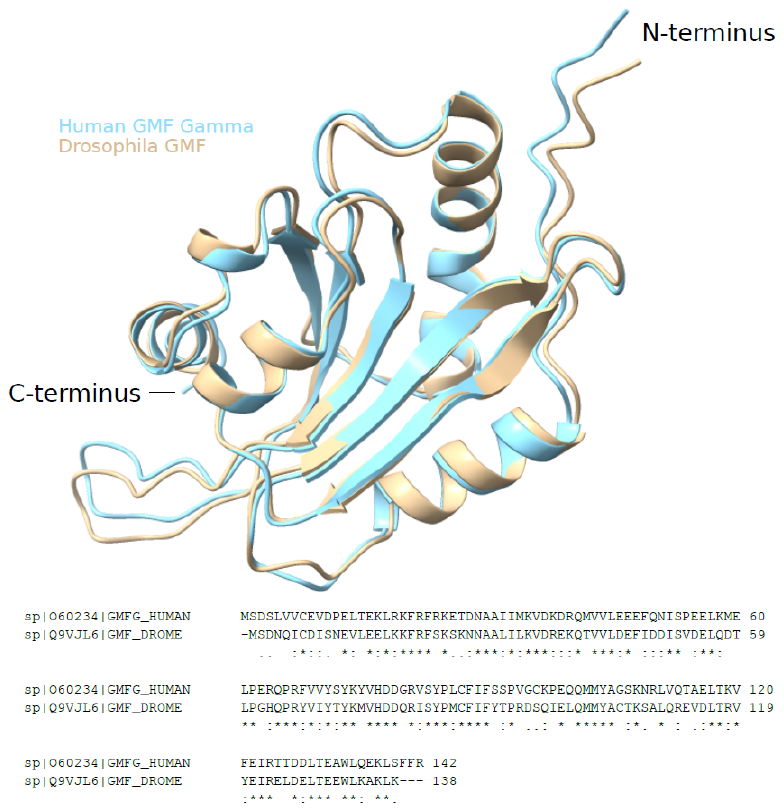

**Fig. S6. Comparing GMF from drosophila and homo sapiens.**

(Top) Aligned Alphafold-generated structures from *Drosophila melanogaster* GMF (Uniprot Q9VJL6, in clay) and *Homo sapiens* GMF Gamma (Uniprot O60234, in light blue). Alignment has been performed using ChimeraX.
(Bottom) Protein sequence alignment using clustalW. Protein similarity is 81%.
